# Epitranscriptomic editing of the RNA N6-methyladenosine modification by dCasRx conjugated methyltransferase and demethylase

**DOI:** 10.1101/2020.10.27.356436

**Authors:** Zhen Xia, Min Tang, Hongyan Zhang, Ryan C. Gimple, Briana C. Prager, Hongzhen Tang, Chongran Sun, Fuyi Liu, Peng Lin, Yutang Mei, Ruoxin Du, Jeremy N. Rich, Qi Xie

**Author notes:** Correspondence (Q.X.).

## Abstract

N6-methyladenosine (m6A) is the most common modification on endogenous RNA transcripts in mammalian cells. Currently, the lack of precise single-nucleotide RNA modification tools makes it challenging to understand the relationship between site-specific RNA methylation and the corresponding phenotypic outcomes. Here, we developed a bidirectional dCasRx epitranscriptome editing platform composed of a nucleus-localized dCasRx conjugated with either a methyltransferase, METTL3, or a demethylase, ALKBH5, to manipulate methylation events at targeted m6A sites in HEK293T and glioma stem cells. This platform specifically and efficiently edited m6A modifications at targeted sites, modulating both gene expression and cell proliferation. We then employed the dCasRx epitranscriptomic editor to further elucidate the molecular function of m6A-binding proteins YTH (DF1, DF2, DF3) family and found that the YTH paralogs act together to mediate RNA degradation. These findings collectively demonstrate that the dCasRx epitranscriptome perturbation platform reported in this study can be employed as site-specific m6A editors for delineating the functional roles of individual m6A modifications in the mammalian epitranscriptome.

## Introduction

Highly dynamic covalent modifications, such as N6-methyladenosine (m6A), pseudouridine (Ψ), 5-methylcytosine (m5C), and N1-methyladenosine (m1A)[1], play critical roles in eukaryotic RNA epitranscriptomic pathways and have been implicated in diseases, especially cancers[2].

Recent studies have demonstrated that m6A is a widespread base modification in the mammalian transcriptome and is enriched near the stop codon and the untranslated regions (UTRs) of mRNAs[3]. m6A signaling modulates RNA secondary structures[4], splicing[5], nuclear localization[6], stability[7, 8], and translation efficiency[9], leading to a wide range of effects on cellular functions. m6A has been shown to regulate many critical processes including the DNA-damage response[10], X-chromosome gene silencing[11], cellular heat shock response[12], hepatic lipid metabolism[13], long-term memory creation[14], spermatogonia differentiation[15], maternal-to-zygotic transition[16], embryonic stem-cell self-renewal and differentiation[17], tumorigenesis[18], and anti-tumor immune responses[19].

The core subunits of the m6A installation(‘writer’) complex in eukaryotic cells, contain methyltransferase-like 3 (METTL3), methyltransferase-like 14 (METTL14)[20], and Wilms tumor 1-associated protein (WTAP) [21]. Within this complex, METTL3 catalyzes a methyl group transfer from S-adenosyl methionine (SAM) to adenine within a single-stranded RNA (ssRNA) sequence motif DRACH (D = A, G, or U; R = A or G; H = A, C, or U), and METTL14 offers RNA binding sites as scaffolds[20]. The m6A eraser complex consisting of AlkB homolog 5 (ALKBH5) and fat mass and obesity-associated protein (FTO), demethylates m6A from the adenine at the same DRACH motif[22, 23]. m6A modifications serve as signals that can be decoded by “reader” proteins, such as YT521-B homology (YTH) families or heterogeneous nuclear ribonucleoprotein families, to impact splicing, stability, translation, and localization of mRNAs[24].

The pleiotropic effects of m6A are mediated by m6A readers in both the nucleus and cytoplasm. Previous studies investigating the impact of m6A on various biological processes and phenotypes have relied on the global manipulation of m6A writers, erasers, and readers, resulting in bulk change of the methylation state of many sites in addition to the regions of interest. As a result, it is difficult to interpret the exact causal relationships between specific m6A modifications and downstream phenotypic changes. For example, writers and erasers that are supposed to have opposite roles in regulating cellular behaviors were found to both maintain survival of glioblastoma (GBM)[25–27]. To elucidate the functional roles of individual m6A modifications in living cells, it is necessary to develop novel m6A editors that modify specific sites of individual transcripts without altering the global RNA methylation pattern.

The development of CRISPR-associated nuclease Cas13 technologies [28] has enabled precise mRNA editing and interrogation of RNA modifications in biological processes. The Cas13 family has an innate ability to bind and cleave ssRNA targeted by a complementary guide RNA[29]. To date, Cas13 effector proteins, including Cas13a (1250 aa), Cas13b (1150 aa), Cas13c (1120 aa), and Cas13d (930 aa), have been reported to show high RNA knockdown efficacy with minimal off-target activity. RfxCas13d (CasRx) is the smallest known enzyme with most substantial targeted knockdown efficiency in the Cas13 family when fused to a nuclear localization sequence (NLS) [30–32]. The CRISPR-Cas9 based editor has been used to manipulate m6A modifications on transcriptomes[33], while CRISPR-Cas13 based platform may become a more promising candidate for m6A editing because it does not require an additional synthetic PAMmer oligonucleotide. Based on previous strategies that fused m6A writers[34] or erasers[35] with Cas13b, we hypothesized that tethering catalytically-inactive CasRx (dCasRx) to m6A writers or erasers could manipulate specific m6A sites targeted by relevant Cas13 guide RNAs. The small sizes of dCasRx epitranscriptome editors allow them to be packaged into lentivirus to study cells that are difficult to transfect using other strategies.

Here, we developed and optimized precise m6A editors through conjugation of dCasRx with METTL3 or ALKBH5, enabling efficient manipulation of individual m6A sites within endo-transcripts and minimal off-target alterations in both normal mammalian cells and cancer cells. While early studies reported that m6A readers YTHDF1, YTHDF2, and YTHDF3 have distinct roles in mediating RNA degradation or translation, a recent study demonstrated that YTHDF1, YTHDF2, and YTHDF3 function together to mediate degradation of m6A-containing mRNAs, thus contradicting the prevailing understanding[36]. Our dCasRx mRNA editing technology enabled us to further interrogate the functional roles of YTH paralogs in cells through controlled manipulation of m6A levels at the YTH paralogs-associated m6A sites and measurement of the alteration of mRNA abundance. Our results demonstrated that high level of methylation levels at YTH paralogs-associated m6A sites enhanced degradation of select endo-transcripts.

## Results

### Designing programmable m6A editors

We fused the dCasRx to an m6A writer, METTL3, or eraser, ALKBH5, to install or remove m6A at targeted sites with single guide RNAs (sgRNAs) (Fig.S1a). Because most methylation and demethylation processing of m6A occurs in the nucleus, we added 2 segments of nuclear localization sequence (NLS) to the dCasRx epitranscriptome editors to promote nuclear localization of the editing complex. The resulting m6A editing complexes are NLS-dCasRx-NLS-METTL3 (dCasRx-M3) and NLS-dCasRx-NLS-ALKBH5 (dCasRx-A5) (Fig.1a, 1b).

**Figure 1.**
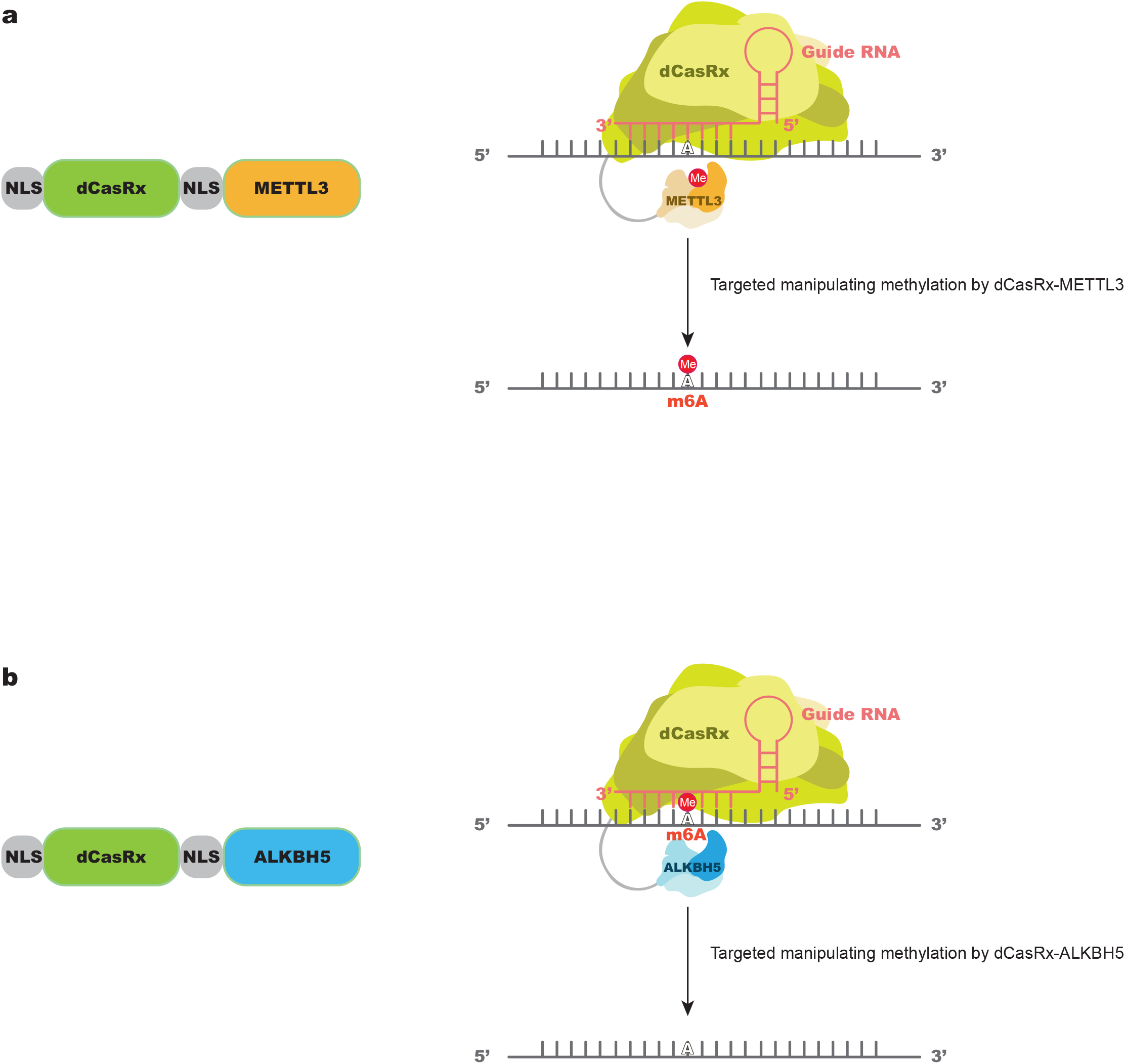
Design of a targeted RNA methylation system. **a**, Proposed strategy for NLS-dCasRx-NLS-METTL3 (dCasRx-M3). dCasRx was fused to METTL3 and nuclear localization sequence (NLS) to mediate site-specific methylation of adenosine to m6A with the presence of sgRNA, as well as to ensure M3 localization in the nucleus. **b**, Proposed strategy for NLS-dCasRx-NLS-ALKBH5 (dCasRx-A5). dCasRx was fused to ALKBH5 and nuclear localization sequence (NLS) to mediate site-specific demethylation of m6A to adenosine with the presence of sgRNA, as well as to ensure dCasRx-A5 localization in the nucleus.

### Validation of editing efficiency of dCasRx epitranscriptome editors in HEK293T cell line

Western blot and confocal imaging were used to confirm that both dCasRx-M3 and dCasRx-A5 were expressed and located in nuclei in the HEK293T cell line (Fig. 2a, 2b). We further generated dCasRx with inactive METTL3 (D395A) or ALKBH5 (H204A) that were also expressed in the nuclei as controls to preclude the possibility that effects were caused by presence of non-specific dCasRx epitranscriptome editors.

**Figure 2.**
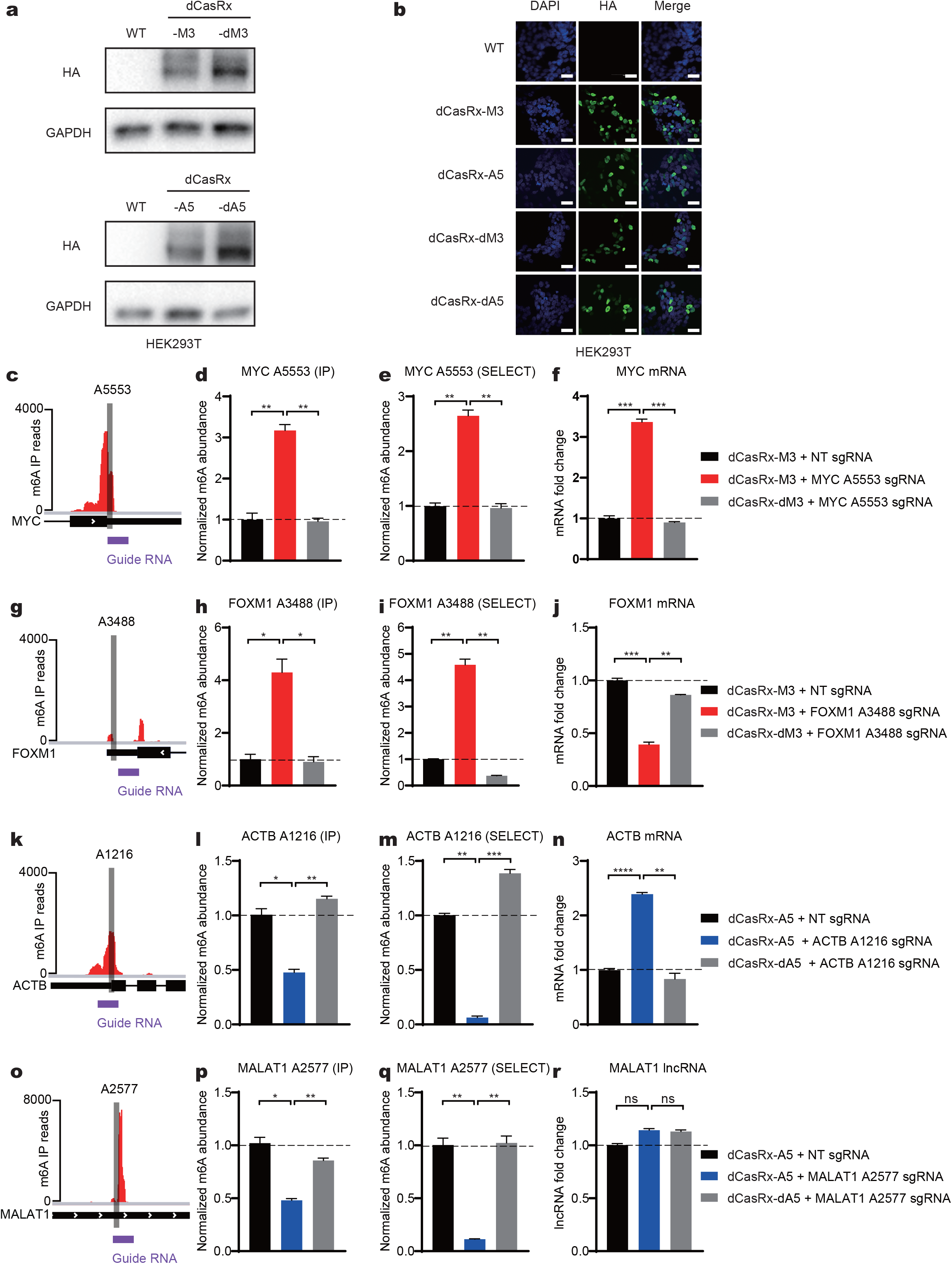
Cellular localization of dCasRx epitranscriptomic editors and targeted manipulation of endogenous transcript methylation in HEK293T cell. **a**, Western blot results of HA-tag demonstrated expression of dCasRx epitranscriptomic editors in living HEK293T cells when treated with dCasRx-M3, dCasRx-A5, dCasRx-dM3, and dCasRx-dA5. **b**, Representative immunofluorescence images of HEK293T cells transfected with HA-tagged dCasRx epitranscriptomic editors (green, HA-tag; blue, DAPI). Scale bars, 40 μm. **c**, Schematic diagram of m6A distribution in MYC mRNA. Grey bar represents dCasRx-M3 or dCasRx-A5 targeted site. Purple bar represents single-strand guide RNA (sgRNA) binding location. **d**, Normalized abundance of m6A at MYC A5553 detected by me-RIP. **e**, Normalized abundance of m6A at MYC A5553 detected by SELECT. **f**, Abundance of MYC mRNA increased after dCasRx-M3 editing in HEK293T cell. **g**, Schematic diagram of m6A distribution in FOXM1 mRNA. **h**, Normalized abundance of altered m6A at FOXM1 A3488 detected by me-RIP. **i**, Normalized abundance of altered m6A at FOXM1 A3488 detected by SELECT. **j**, Abundance of FOXM1 mRNA decreased after dCasRx-M3 editing in HEK293T cell. **k**, Schematic diagram of m6A distribution in ACTB mRNA. **l**, Normalized abundance of m6A at ACTB A1216 detected by me-RIP. **m**, Normalized abundance of m6A at ACTB A1216 detected by SELECT. **n**, Abundance of ACTB mRNA increased after dCasRx-A5 editing in HEK293T cell. **o**, Schematic diagram of m6A distribution in MALAT1 RNA. **p**, Normalized abundance of altered m6A at MALAT1 A2577 detected by me-RIP. **q**, Normalized abundance of altered m6A at MALAT1 A2577 detected by SELECT. **r**, Abundance of MALAT1 lncRNA has not changed after dCasRx-A5 editing in HEK293T cell. Data are displayed as mean ± SEM (ANOVA, ns: not significant, *:P < 0.05, **:P < 0.01, ***:P < 0.001, ****:P < 0.0001, n=3).

To fully characterize the editing window of dCasRx-M3 or dCasRx-A5, we designed 9 different sgRNAs surrounding the A1216 site of the ACTB transcript and co-transfected these sgRNAs along with dCasRx-M3 or dCasRx-A5 into HEK293T cells. sgRNAs were tiled across a 20bp region spanning the targeted site with a 3nt gap between each sgRNA, and one additional sgRNA covered the targeted site from −1 to +1 nt (Fig. S1b). dCasRx epitranscriptomic editors demonstrated good editing efficiency within the editing window from −4 to +1 nt relative to the targeted site (Fig. S1c-S1d). In addition, dCasRx epitranscriptomic editors also exhibited a high editing efficiency using the +7 nt sgRNA.

Traditional methods such as methylated RNA immunoprecipitation (me-RIP) coupled with RT–qPCR or a sequencing-based approach, methylation-individual-nucleotide resolution Cross-Linking and Immunoprecipitation (miCLIP), lack either single-nucleotide resolution or quantitative features. To circumvent these limitations, we used an established single-base elongation- and ligation-based qPCR amplification method, termed SELECT, to measure m6A level alterations[37]. The SELECT technique employs the ability of m6A to hinder the elongation activity of DNA polymerase and the ability of DNA-ligase to selectively catalyze nick ligation between the elongated Up Probe and Down Probe. The final products were quantified by qPCR to reflect the m6A abundance. Using an orthogonal method, we verified that the m6A alteration detected using SELECT reliably reflected methylation level changes at targeted sites, as described by previous studies[38].

We next demonstrated that the nucleus-localized dCasRx epitranscriptomic editors could install or remove m6A modifications on endogenous transcripts in HEK293T cells. We first evaluated the ‘writing’ efficiency of dCasRx-M3 using adenine sites that have a low degree of methylation in native HEK293T cells. Adenine A5553 within the 3′-UTR of MYC mRNA (Fig. 2c) and adenine A3488 within the 3′-UTR of FOXM1 mRNA (Fig. 2g) were targeted. With co-transduction of the targeting sgRNA, dCasRx-M3 expression increased MYC A5553 methylation (Fig. 2d, 2e) followed by an upregulation of MYC mRNA (Fig. 2f). Transduction with a different sgRNA increased FOXM1 A3488 methylation followed by a downregulation of FOXM1 mRNA (Fig. 2j). These results indicated that specific m6A sites are editable by dCasRx epitranscriptomic editors. In contrast, transduction with methyltransferase-inactive variants (dCasRx-dM3) of these constructs at comparable expression to their active counterparts did not increase methylation level of target sites in the corresponding mRNAs. Another control using dCasRx-M3 with non-targeted sgRNA (NT) also showed no significant alteration of m6A levels at targeted sites, confirming that expression of dCasRx epitranscriptomic editors alone does not induce m6A changes at targeted sites. We then evaluated the ‘erasing’ efficiency of dCasRx-A5 using adenine sites that have a high degree of methylation in native HEK293T cells. Adenine A1216 within the 3′-UTR of ACTB mRNA (Fig. 2k) and adenine A2577 of MALAT1 mRNA (Fig. 2o) were targeted. Both SELECT and me-RIP showed a significant decrease of methylation at targeted sites in the dCasRx-A5 edited cells compared with the NT or dCasRx-dA5 group (Fig. 2l, 2m, 2p, 2q), indicating that dCasRx-A5 is able to remove m6A modifications, which induced an increase of ACTB mRNA (Fig. 2n) but had no effect on MALAT long non-coding RNA (Fig. 2r).

To evaluate the off-target activity of dCasRx epitranscriptomic editors, we also measured changes in m6A levels in three other adenine sites in HEK293T cells. The levels of m6A at non-targeted adenine sites were not affected in either dCasRx-M3 groups (Fig. S2a-S2c, S2g-S2i) or dCasRx-A5 groups (Fig. S2d-S2f, S2j-S2l), suggesting low off-target editing activity of the dCasRx epitranscriptomic editors.

### Manipulating m6A levels at FOXM1 and MYC mRNA regulates GSC proliferation

After validating the editing efficiency of dCasRx epitranscriptomic editors in HEK293T cells, we applied the dCasRx epitranscriptomic editors to glioblastoma stem cells (GSCs). Unlike HEK293T cells, GSCs could not be easily transfected with plasmids carrying dCasRx-M3 or dCasRx-A5. Thus, we developed a lentiviral delivery approach to employ dCasRx editors in GSCs. GSCs that expressed dCasRx epitranscriptomic editors were enriched by selective culturing in a puromycin-containing medium, owing to its resistance to puromycin. Both the dCasRx-M3 and dCasRx-A5 constructs were expressed and localized in nuclei of both GSC 3565 and GSC 468 cells (Fig. 3a, 3b). The sgRNAs were also packaged into lentiviruses using the same procedure. GSCs expressing dCasRx-M3 or dCasRx-A5 were then infected with these sgRNA-lentiviruses.

**Figure 3.**
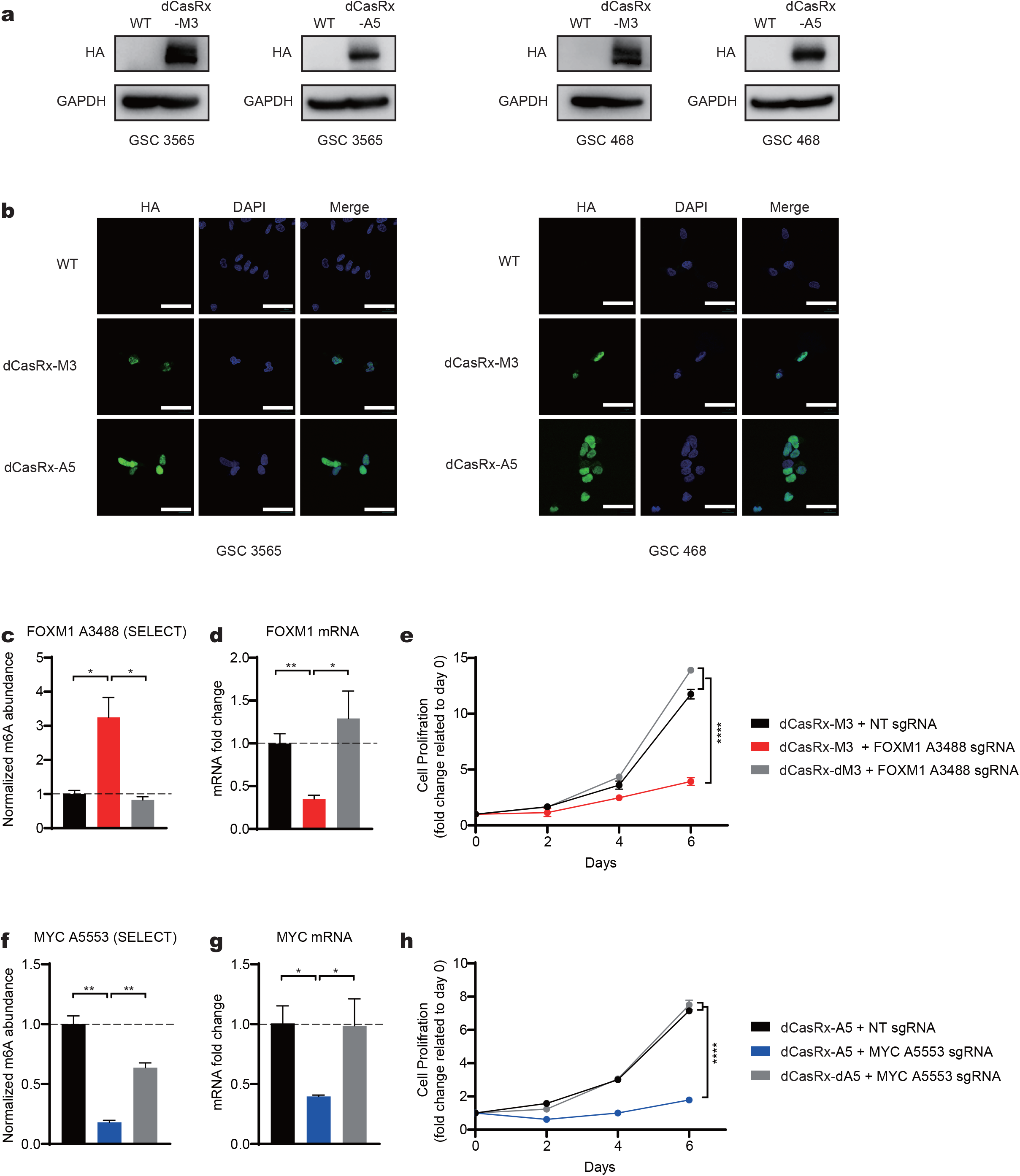
Targeted manipulation of m6A level via dCasRx epitranscriptomic editors on endogenous transcripts affects GSC proliferation. **a**, Expression of dCasRx epitranscriptomic editors dCasRx-M3 or dCasRx-A5 in GSC 3565 and GSC 468. **b**, Representative immunofluorescence images of GSCs transfected with HA-tagged dCasRx epitranscriptomic editors (green, HA-tag; blue, DAPI). Scale bars, 40 μm. **c**, Methylation at A3488 of endogenous mRNA FOXM1 in GSC 3565 increased by 3 folds by dCasRx-M3 editing. **d**, mRNA expression level of FOXM1 decreased after dCasRx-M3 editing in GSC 3565. **e**, Proliferation of GSC 3565 decreased with an increased m6A level at FOXM1 A3488 mediated by dCasRx-M3, compared to NT group and dCasRx-dM3 group. **f**, A decrease of methylation at A5553 of endogenous mRNA MYC was mediated by dCasRx-A5 editing in GSC 3565. **g**, mRNA expression level of MYC decreased after dCasRx-A5 editing in GSC 3565. **h**, Proliferation of GSC 3565 decreased with a decreased m6A level at MYC A5553 mediated by dCasRx-A5, compared to NT group and dCasRx-dA5 group. Data is represented as mean ± SEM. (ANOVA, *: P value < 0.05; **: P value < 0.01; ***: P value < 0.001; ****: P value < 0.0001, n = 3).

The transcription factors FOXM1 and MYC are known to play essential roles in regulating GSC proliferation, self-renewal, and tumorigenicity. The mRNAs of FOXM1 and MYC also contain adenine sites that can be methylated by m6A. Our programmable epitranscriptome writer dCasRx-M3 increased the m6A level at the A3488 site of the FOXM1 transcript (Fig. 3c); and the programmable eraser dCasRx-A5 reduced the m6A level at the A5553 site of the MYC transcript in GSC 3565 cells compared to non-targeting and catalytically-inactive controls (Fig. 3f).

Low m6A signal at the 3’UTR on FOXM1 mRNA enhances the expression of FOXM1 nascent transcripts [26], whereas m6A at the 3’UTR in MYC stabilizes its mRNA[39, 40]. Consistent with these reported findings, targeted m6A methylation at the FOXM1 transcript A3488 site using dCasRx-M3 led to a decrease of FOXM1 mRNA (Fig. 3d). Additionally, targeted m6A demethylation at the MYC transcript A5553 site downregulated the expression of MYC mRNA (Fig. 3g). Both NT sgRNA and catalytically inactive constructs were used as controls. Downregulation of FOXM1 or MYC mRNA induced by dCasRx editing significantly restricted the proliferation of GSC 3565 (Fig. 3e, 3h). These results demonstrated that our dCasRx epitranscriptomic editors could manipulate m6A levels on endo-transcripts in GSCs and subsequently regulate GSC proliferation.

### YT521-B homology Domain-containing Family (YTHDF) proteins mediate degradation of m6A-mRNAs

The effects of m6A in cytosolic transcripts are mediated by a complex network of interactions between specific m6A sites and specific members of the YTHDF of m6A-binding proteins. The YTHDF family includes three paralogs, YTHDF1 (DF1), YTHDF2 (DF2), and YTHDF3 (DF3), each with distinct reported functions. In the classical model, DF1 enhances mRNA translation, DF2 promotes mRNA degradation, and DF3 works as a co-factor with DF1 or DF2 to enhance either mRNA translation or degradation [24]. However, a recent study found that the m6A sites binding with DF1, DF2, or DF3 are highly similar, and all three paralogs function together to mediate degradation of m6A-tagged mRNAs[36].

To resolve this controversial function of the YTHDF proteins, we analyzed a database (GSE78030) which presented the distribution of YTHDF paralogs in endo-transcriptome in HEK293T cells[11] and screened mRNA candidates that contained m6A sites for YTHDF paralogs binding. Among these, m6A sites of SQLE A0724 (Fig. 4a) and ACTB A1216 (Fig. S3a) presented high-affinity binding sites to all three YTHDF paralogs. The m6A site of SQLE A0724 was located at the 5’UTR, which has potential to bind eIF4F to start cap-independent translation[9], while the m6A site of ACTB A1216 was located at the 3’UTR and near the stop codon. To examine whether a single site binding with all three DFs in the 3′ UTR or 5’UTR contributed to the degradation of relevant mRNAs, we designed sgRNAs for the corresponding targeted sites. For both SQLE (Fig. 4b, 4c) and ACTB (Fig. S4b, S4c), a significant decrease of mRNA expression was induced by m6A installation via dCasRx-M3 editing (panels). Conversely, a significant increase of relevant mRNAs was induced by m6A removal via dCasRx-A5 editing (Fig. 4e, 4f, 2m, 2n). To confirm that the effects of m6A editing were mediated by changes in degradation, we assayed mRNA stability. Consistent with our findings, m6A removal led to accelerated mRNA degradation and thus decreased mRNA levels (Fig. 4d, 4g, S3d, S2e).

**Figure 4.**
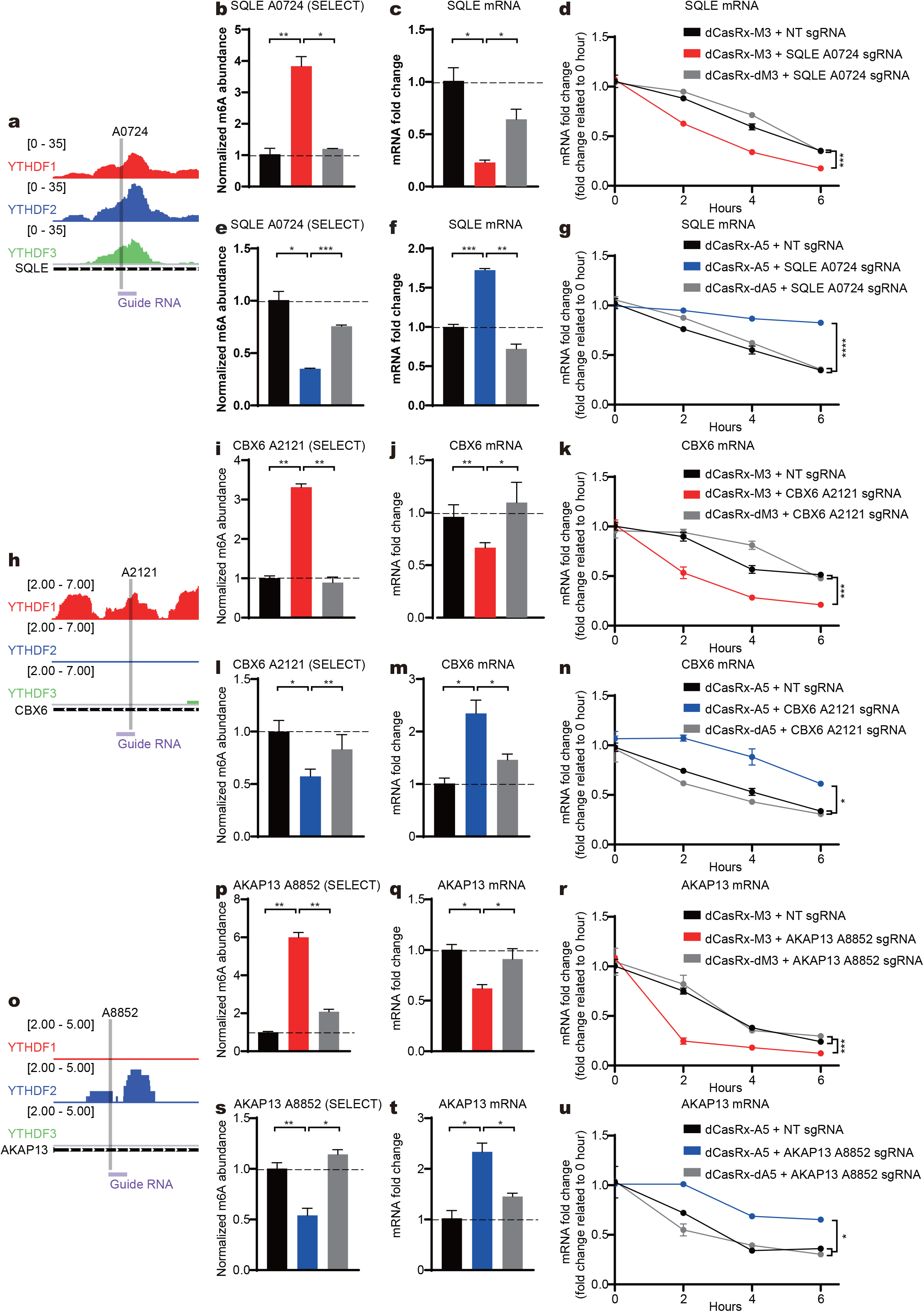
m6A sites binding with DF paralogs control the degradation of endogenous transcripts in HEK293T cells. **a**, **h**, **o**, Schematic diagrams of distribution of DF paralogs, DF1 (red), DF2 (blue), and DF3 (green), in endogenous SQLE mRNA (**a**), CBX6 mRNA (**h**), AKAP13 mRNA (**o**). Grey bars represent dCasRx-M3 or dCasRx-A5 targeted sites. Purple bars represent sgRNA binding location. **b**, **i**, **p**, Normalized abundance of altered m6A at SQLE A0724 (**b**), CBX6 A2121 (**i**), AKAP13 A8852 (**p**) edited by dCasRx-M3. **c**, **j**, **q**, Abundance of SQLE mRNA (**c**), CBX6 mRNA (**j**), AKAP13 mRNA (**q**) decreased after dCasRx-M3 editing. **d**, **k**, **r**, mRNA degradation measurement of SQLE (**d**), CBX6 (**k**), AKAP13 (**r**) in HEK293T cells edited with dCasRx-M3. **e**, **l**, **s**, Normalized abundance of altered m6A at SQLE A0724 (**e**), CBX6 A2121 (**l**), AKAP13 A8852 (**s**) edited by dCasRx-A5. **f**, **m**, **t**, Abundance of SQLE mRNA (**f**), CBX6 mRNA (**m**), AKAP13 mRNA (**t**) increased after dCasRx-A5 editing. **g**, **n**, **u**, mRNA degradation measurement of SQLE (**g**), CBX6 (**n**), AKAP13 (**u**) in HEK293T cells transfected with dCasRx-A5. Data is represented as mean ± SEM. (ANOVA, *: P value < 0.05; **: P value < 0.01; ***: P value < 0.001; ****: P value < 0.0001. n = 3).

To further verify the individual functional roles of m6A sites interacting with DF1 or DF2, we identified two gene candidates containing m6A sites for each DF and designed sgRNAs accordingly. By writing at m6A sites that bind DF1 (Fig. 4h, S3f), we demonstrated that increased level of m6A at CBX6 A2121 (Fig. 4i, 4l) and Sbrbp1 A3240 (Fig. S3g, S3i) led to downregulation of corresponding mRNA (Fig. 4j-4k, 4m-4n, S3h, S3j). By regulating m6A sites dominantly binding with DF2, we showed that increased level of m6A at AKAP13 A8852 (Fig. 4o-4p, 4s) and MLLT3 A1068 (Fig. S3k-S3l, S3n) also led to downregulation of corresponding mRNAs (Fig. 4q, 4t, S3m, S3o), suggesting that the installation of m6A at those sites enhanced DF2-mediated mRNA degradation (Fig. 4r, 4u). The observed regulatory effects of m6A sites interacting with DF1 contradicted the classical model. Thus we sought to investigate whether inclusion of DF3 would induce different alterations. We next screened two m6A sites dominantly binding with both DF1 and DF3 but not with DF2, GSTP1 A0659 (Fig. S4a) and HDGF A1908 (Fig. S4h), and used dCasRx-M3 or dCasRx-A5 to manipulate their methylation level. The results demonstrated that increased methylation level at targeted sites binding both DF1 and DF3 still promoted mRNA degradation (Fig S4b-S4g, S4i-l).

By modulating the methylation level of targeted sites at YTHDF paralogs with dCasRx-M3 and dCasRx-A5, we found that the m6A sites binding with DF1, DF2, and DF3 all resulted in m6A-mediated mRNA degradation, thus supporting the recent findings[36] rather than the classical model[24].

## Discussion

The advancement of various gene editors enables functional analysis of epigenomic marks and facilitates investigation of epigenetic control over biological processes [41, 42]. RNA modifications are essential parts of the epitranscriptome, and the highly dynamic and reversible m6A modifications in mammalian transcriptomes regulate nearly all aspects of RNA metabolism and functionality[43, 44]. However, much of our current knowledge about m6A was mainly based on genetic perturbations induced by global overexpression or knockout of relevant genes, which modify the whole transcriptome instead of target sites of interest. To interrogate the site-specific effects of m6A interacting with multiple readers, a strategy to manipulate targeted m6A sites within endo-transcripts is essential.

In this work, we developed the first dCas13Rx-mediated epitranscriptomic editors that coupled with RNA methyltransferase METTL3 or demethylase ALKBH5 to achieve bidirectional modulation of targeted m6A sites in living cells. We demonstrated that nucleus-located dCasRx epitranscriptomic editors enabled site-specific m6A installation or removal with low off-target alterations. In addition to being transfected into normal human cells, the dCasRx epitranscriptomic editors could be packaged into lentiviruses, owing to their small sizes, and be used to investigate the function of single m6A sites in glioblastoma cells that were difficult to be transfected by other means. We demonstrated that m6A played opposite roles in the regulation of FOXM1 and MYC mRNA – decreased expression of FOXM1 transcripts but increased MYC expression. Alterations in the expression level of FOXM1 and MYC subsequently affected GSCs proliferation. Lastly, using the dCasRx epitranscriptomic editors, we demonstrated that increased methylation of YTH paralogs-bound sites induced mRNA degradation, in contrast to the described function in the classical model. Collectively, these results demonstrate that our dCasRx system can be used to dissect previously unclear interactions and to elucidate the causal relationships between m6A modifications and phenotypes.

This work demonstrates a proof-of-concept dCasRx-based strategy with the potential for high-throughput screening of m6A modifications in whole epitranscriptome using a suitable sgRNA library. Judging from the success of high-throughput functional genomic screening based on CRISPR-Cas9 technology[45] and verified efficiency of dCasRx epitranscriptomic editors, our technology has broad applications in a variety of studies including investigation of epitranscriptome of difficult-to-transfect cancer cells.

## MATERIALS AND METHODS

### Design of dCasRx epitranscriptomic editors

dCasRx epitranscriptomic editors were constructed by fusing candidate methyltransferase METTL3 (Δ458-506) or demethylase ALKBH5 to the C terminus of SV40 NLS-dCasRx (Addgene plasmid # 118634) via a nucleoplasmin NLS (K-R-P-A-A-T-K-K-A-G-Q-A-K-K-K-K)-G-S-S linker. The dCasRx-dMETTL3 conjugate contained a single mutation at D395A to form the dCasRx-dM3. The dCasRx-dALKBH5 conjugate contained a single mutation at H204A to form the dCasRx-dA5. A Flag (D-Y-K-D-D-D-D-K)-G-G-G-G-G-HA (Y-P-Y-D-V-P-D-Y-A) signal was added at the C terminus of all epitranscriptomic editors as a tag for detection.

### Design of sgRNAs

Information of distributions of DF paralogs was based on a database GSE78030[11]. We selected adenine sites that have been reported accessible to methylation modifications by m6A in a database GSE63753 [46] as target sites for designing sgRNAs. Designed sgRNAs were checked using the NCBI BLAST (https://blast.ncbi.nlm.nih.gov/Blast.cgi) to avoid unwanted mRNA off-target bindings in the human genome. Sequence of sgRNAs are provided in Supplementary Table S1.

### Cell culture

HEK293T cells were cultured in Dulbecco’s Modified Eagle Medium (DMEM, Gibco, # C11995500CP) supplemented with 10% fetal bovine serum (FBS, Gibco, # 10099-141C), 1% penicillin/streptomycin (Hyclone # SV30010), and 1% GlutaMax™ Supplement (Gibco, # 35050-061) at 37°C, 5% CO2. GSC 3565 and GSC 468 cells were cultured in Neurobasal™-A Medium (Gibco, # 12349-015) supplemented with 1% B-27™ supplement (Gibco, # 12587-010), 20 ng/ml recombinant human EGF protein (R&D, # 236-EG), 20 ng/ml recombinant human FGF basic protein (R&D, # 4114-TC),1% penicillin/streptomycin (P/S, Invitrogen, SV30010), 1% sodium pyruvate (Gibco, # 11360-070), and 1% GlutaMax™ supplement (Gibco, # 35050-061) at 37°C, 5% CO2.

### Plasmid transfection

Plasmid transfection was carried out using LipoD293™ In Vitro DNA Transfection Reagent (SignaGen Laboratories, SL100668) following manufacture's protocol. For six-well assays, HEK293T cells were co-transfected with 1.5 μg dCasRx conjugate plasmid and 1.5 μg sgRNA plasmid per well. Transfected cells were cultured under normal conditions (37 C, 5% CO2) for 36 hours before analysis.

### Lentivirus packaging

In 100 mm dishes, HEK293T cells were transfected using 6 μg of the proposed plasmid, 4 μg of psPAX2 (Addgene plasmid # 12260), and 2 μg of pMD2.G plasmid (Addgene plasmid # 12259) per well with LipoD293™ In Vitro DNA Transfection Reagent. The transfected cells were incubated under normal conditions for 48 hours and then harvested. The supernatant of HEK293T cell culture was collected after centrifugation at room temperature (RT). Concentration of virus was quantified using Lentivirus Concentration Solution (Genomeditech, # GM-040801-100) following manufacture's protocol. The lentivirus was stored at −80°C before use.

### Constructing GSCs expressing dCasRx epitranscriptomic editors and sgRNAs

To generate dCasRx epitranscriptomic editors edited GSCs, GSCs were first digested as single cells followed by incubation with dCasRx epitranscriptomic editors lentiviruses for 12 hours in culture medium. GSCs expressing dCasRx epitranscriptomic editors were selected in medium containing Puromycin Dihydrochloride (Beyotime, # ST551-10mg) for 5 days, owing to the puromycin-resistance of dCasRx conjugate plasmids. Then, GSCs expressing dCasRx epitranscriptomic editors were infected with sgRNAs lentiviruses with Blasticidin S HCl (Beyotime, # ST018-1ml) resistance. GSCs were selected with Blasticidin S HCl supplemented medium for 3 days to select GSC cells expressing both dCasRx and sgRNAs. The engineered GSCs were used for following experiments.

### Protein extraction and western blotting

Cells were washed twice with PBS and lysed with RIPA Lysis Buffer (Beyotime, # P0013C) supplemented with phenylmethanesulfonyl fluoride (Sigma-Aldrich, # p7626-1g) and cOmplete™(Roche, # 4693132001) on ice. Supernatant of cell lysates was collected after centrifugation and denatured at 100°C with Loading buffer. Samples were loaded onto an 8% 15-well SDS-PAGE gel, and the gel was transferred to a 0.45-μm polyvinylidene difluoride membrane (Merckmillipore, # IPVH00010) after electrophoresis. Membranes were blocked with NON-Fat Powdered Milk (Sangon Biotech, # A600669-0250) and stained with HA-Tag (C29F4) rabbit mAb (Cell Signaling Technology, # 3724S) and GAPDH monoclonal antibody (Proteintech, # 60004-1-Ig) in TBST (TBS + 0.5% Tween-20) with 1% Bovine serum albumin (BSA, Sangon Biotech, # A600332-0100) overnight at 4 °C. After washing three times with TBST, membranes were incubated with secondary antibodies, anti-rabbit HRP-linked antibody (Cell Signaling Technology, # 7074) and anti-mouse HRP-linked antibody (Cell Signaling Technology, # 7076), with 5% nonfat dry milk in TBST for 1 hour at room temperature. The membrane was then washed three times with TBST and imaged using ChemiDoc XRS+ System (BIO RAD, # 1708265).

### Immunofluorescence microscopy

An HA epitope tag (Y-P-Y-D-V-P-D-Y-A) was cloned onto the C terminus of dCasRx epitranscriptomic editors for detecting the location of dCasRx epitranscriptomic editors. HEK293T and GSC cells expressing dCasRx epitranscriptomic editors were seeded and cultured on coverglasses (Solarbio, # YA0350) in 24-well plates. After 36 hours of incubation, the culture medium was discarded, and the coverglasses were washed once with PBS gently. Cells were fixed with 4% paraformaldehyde (PFA, Servicebio, # G1101) for 10 minutes. Cells were then washed three times with PBS and permeabilized with PBS + 0.2% Triton-X100 (PBST) for 15 min at room temperature. Cells were blocked in blocking buffer (10% BSA in PBST) for 30 min and stained with HA-Tag (C29F4) rabbit mAb in blocking buffer overnight at 4°C. Cells were then washed three times with PBST and stained with donkey anti-rabbit IgG (H+L) Alexa Fluor Plus 488 (Thermo Fisher Scientific, # A32790) in blocking buffer for 1 hour at room temperature followed by 1 hour of DAPI (Roche, # 33495822) staining. Images were acquired using a confocal laser scanning microscope (Olympus, FV3000-IX83).

### RNA isolation and RT-qPCR

For RNA extraction, fresh cells were first washed twice with PBS, and then total RNA was extracted using TRIzol™ Reagent (Thermo Fisher Scientific, # 15596018) following manufacturer’s protocol. For each sample, 1 μg of total RNA was used for reverse transcription to cDNA using Novoscript Plus All in one 1st Strand cDNA Synthesis SuperMix (Novoprotein, # E047-01S). The cDNA templates were used in RT-qPCR quantification. Each 20 μl qPCR reaction contained 1 μl of cDNA, 1 μM forward and reverse primers, and 10μl of 2X SYBR Green Master Mix (Novoprotein, # E096-01S). Reaction mixture was heated at 95 °C for 1 minute, followed by 40 repeated cycles with the following conditions: 95 °C for 20 seconds and 60 °C for 60 seconds. All assays were repeated with three independent experiments. The primers used in RT-qPCR assays are listed in the Supplementary Table S2.

### SELECT technology for detection of m6A

Detection of m6A at targeted sites was based on the SELECT technology modified from a previous protocol[37]. For each sample, 1 μg of total RNA was incubated with 40 nM Up primer, 40 nM Down primer, and 5 μM dNTP (New England Biolabs, # N0446S) in 17.5 μl 1x CutSmart buffer (New England Biolabs, # B7204S). A progressive annealing cycle was carried out: 1 minute each at 90 °C, 80 °C, 70 °C, 60 °C, 50 °C, and 40 °C for 6 minutes. Subsequently, 2.5 μl of enzyme mixture containing 0.01 U Bst 2.0 DNA polymerase (New England Biolabs, M0537S), 0.5 U SplintR ligase (New England Biolabs, # M0375S), and 10 nmol ATP (New England Biolabs, # P0756S) were added to the 17.5 μl annealing products. The final 20 μl reaction mixtures were incubated at 40 °C for 20 minutes, denatured at 80 °C for 20 minutes, and then kept at 4 °C. Afterward, 2 μl of final products were transferred to a reaction mixture containing 200 nM SELECT common primers and 2X SYBR Green Master Mix for qPCR analysis. The run cycle was set up as: 95 °C for 1 min, followed by 40 cycles of (95 °C, 20 seconds; 60 °C, 60 seconds). The SELECT products of targeted sites were normalized to the RNA abundance of corresponding transcripts containing m6A sites. All assays were performed with three independent experiments. Primers used in the SELECT assays are listed in Supplementary Table S3.

### Low input m6A-RIP coupled with qPCR

The strategy of low input m6A-RIP was modified from Zeng’s protocol[38]. Total RNA from cells was extracted using Trizol reagent, and DNA was removed by adding Dnase I (New England Biolabs, # M0303S). A total volume of 5 μg total RNA was resuspended to 18 μl with RNase-free water. 2 μl of 10X RNA Fragmentation Buffer (100 mM Tris-HCl, 100 mM ZnCl2 in nuclease-free H2O) was added to the RNA and incubated in a preheated thermal cycler for 5 minutes at 95°C for fragmentation. The total RNA was chemically fragmented into ~200-nt-long fragments. To prepare antibody-bead for me-RNA immunoprecipitation, 15 μl of protein-A magnetic beads (Thermo Fisher Scientific, # 10002D) and 15 μl of protein-G magnetic beads (Thermo Fisher Scientific, # 10004D) were incubated with 5 μg anti-m6A antibody (Cell Signaling Technology, D9D9W) at 4°C overnight. The prepared antibody-bead mixture was resuspended in 500 μl of IP reaction mixture containing fragmented total RNA and incubated for 2 hours at 4°C. The RNA reaction mixture was then washed twice in 1000 μl of IP buffer, followed by elution buffer with continuous shaking for 1 h at 4°C. Additional phenol-chloroform isolation and ethanol precipitation treatments were performed to purify the RNA. We also used NovoProtein Novoscript Plus All in one 1st Strand cDNA Synthesis SuperMix followed by qPCR for quantification. All assays were performed with three independent experiments. Primers used in the low-input m6A-RIP assay are listed in Supplementary Table S4.

### Cell proliferation assay

2000 GSC cells were seeded into 96-well plate well with 200 μl medium at the beginning. Before detection, cells were cooled down to RT, followed by adding 40 μl of Cell Titer (Promega, # G7572) per well. The plate was shaken at 120 r.p.m. for 15 mins at RT, and 150 μl of the final product was used for proliferation detection on a microplate reader (Thermo, Varioskan LUX). All assays were performed with three independent experiments.

### mRNA stability assay

HEK293T cells co-transfected with dCasRx conjugate and relevant sgRNA vectors were treated with 5 μg/ml transcription inhibitor ActinoMYCin D (Selleck, # S8964) and collected at various time points (0, 1, 2, and 6 hours). The total RNA was extracted and reverse transcribed as described before. The cDNA templates were used in RT-qPCR quantification. The primers used in RT-qPCR assay are listed in Supplementary Table S2. All assays were performed with three independent experiments.

### Quantification and Statistical Analyses

All statistical analyses are described in the figure legends. Two-way repeated-measures ANOVA was used for statistical analysis with Dunnett multiple hypothesis test correction.

## Data Availability

All data accessed from external sources and prior publications have been referenced in the text. Additional data will be made available upon request.

## Acknowledgments

This work is supported by the National Natural Science Foundation of China (82073268) and Westlake Education Foundation.

## Contact for Reagent and Resource Sharing

Further information and requests for resources and reagents should go to the lead contact Q.X.

---

**Figure S1.**
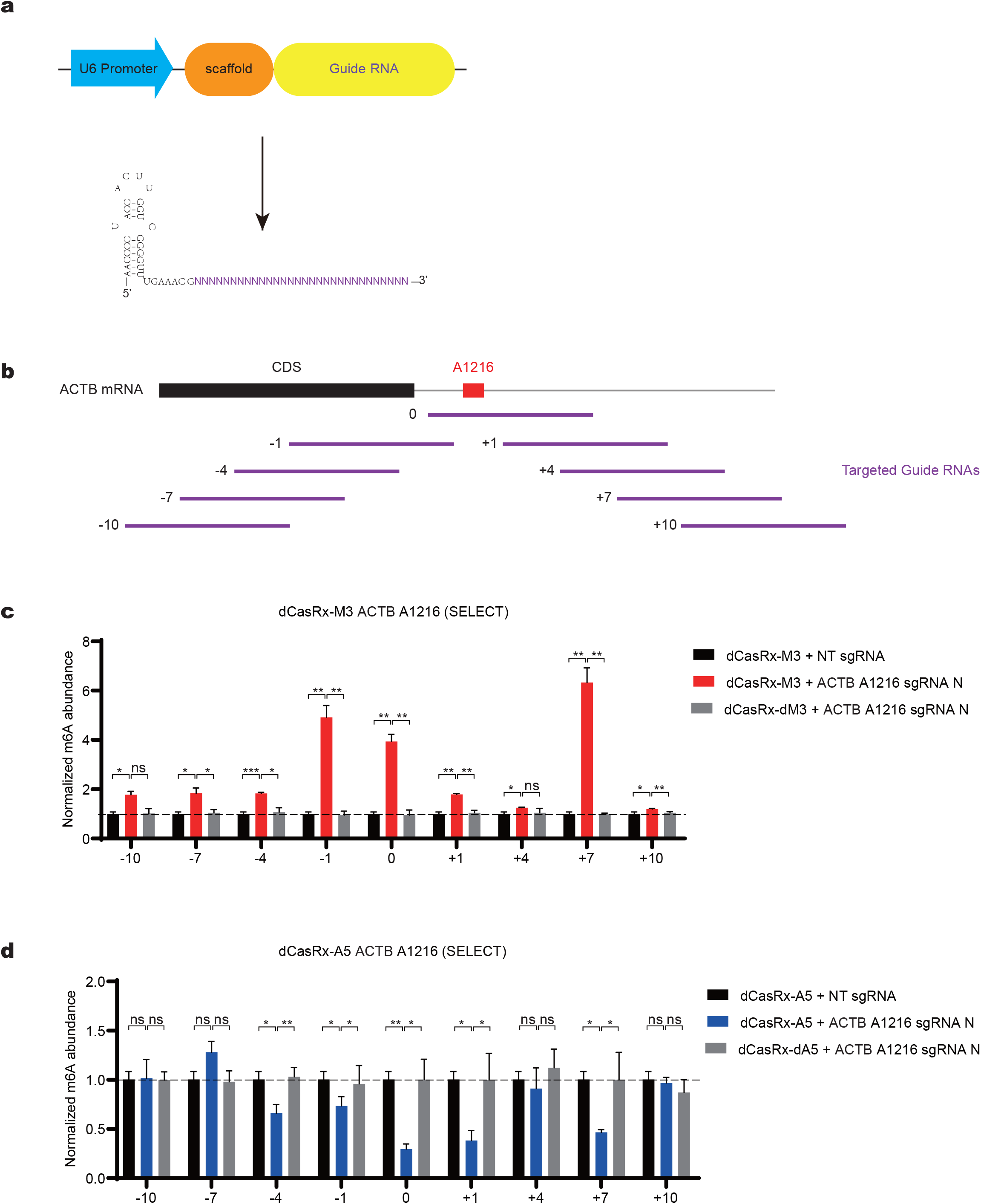
dCasRx epitranscriptomic editors editing window on ACTB mRNA. **a**, Schematic diagram of sgRNA designed for dCasRx epitranscriptomic editors. **b**, Illustration of the sgRNAs designed for targeting ACTB A1216. each 30-nt sgRNAs (purple) ending (−1, −4, −7, −10) or starting (+1, +4, +7, +10) at the indicated bp from the targeted site. **c**, Normalized abundance of m6A altered by dCasRx-M3 with sgRNAs targeting ACTB A1216. **d**, Normalized abundance of m6A altered by dCasRx-A5 with sgRNAs targeting ACTB A1216. Data is represented as mean ± SEM. (ANOVA, *: P value < 0.05; **: P value < 0.01; ***: P value < 0.001,n = 3).

**Figure S2.**
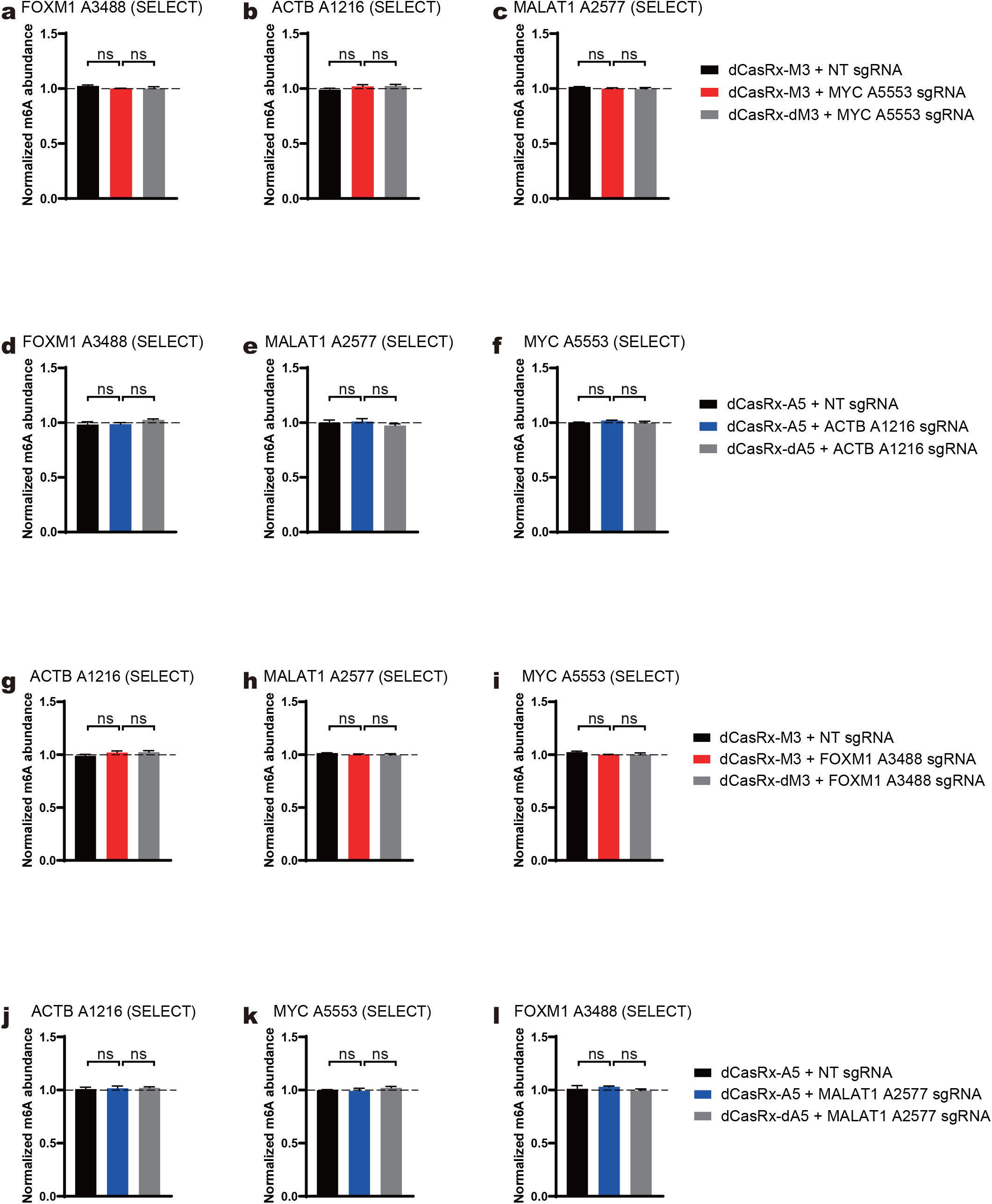
m6A level at non-targeted adenine sites in HEK293T cells. **a-c**, Normalized abundance of altered m6A at non-targeted adenine sites in FOXM1 A3488 (**a**), ACTB A1216 (**b**), and MALAT1 A2577 (**c**) using sgRNA for MYC A5553, detected by SELECT. **d-f**, Normalized abundance of altered m6A at non-targeted adenine sites in FOXM1 A3488 (**d**), MALAT1 A2577 (**e**), and MYC A5553 (**f**) using sgRNA for ACTB A1216 detected by SELECT. **g-i**, Normalized abundance of altered m6A at non-targeted adenine sites in ACTB A1216 (**g**), MALAT1 A2577 (**h**), and MYC A5553 (**i**) using sgRNA for FOXM1 A3488 detected by SELECT. **j-l**, Normalized abundance of altered m6A at non-targeted adenine sites in ACTB A1216 (**j**) MYC A5553 (**k**), and FOXM1 A3488 (**l**) using sgRNA for MALAT1 A2577 detected by SELECT. Data is represented as mean ± SEM. (ANOVA, ns: not significant, n = 3).

**Figure S3.**
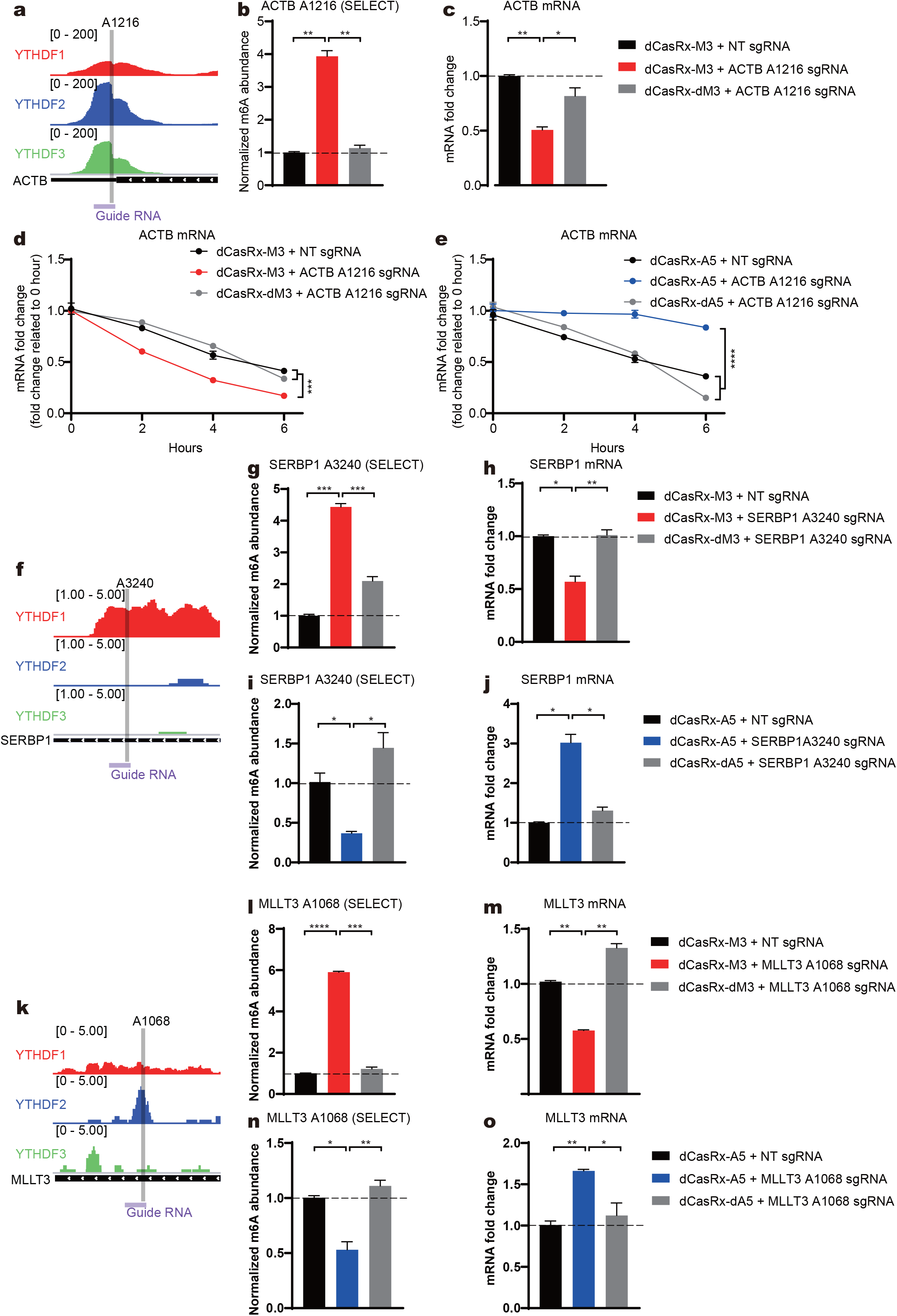
m6A sites binding with DF paralogs control the degradation of endogenous transcripts in HEK293T cells. **a**, **f**, **k**, Schematic diagrams of distribution of DF paralogs, DF1 (red), DF2 (blue), and DF3 (green), in endogenous ACTB mRNA (**a**), SERBP1 mRNA (**f**), MLLT3 mRNA (**k**). Grey bars represent dCasRx-M3 or dCasRx-A5 targeted sites. Purple bars represent sgRNA binding location. **b**, **g**, **l**, Normalized abundance of altered m6A at ACTB A1216 (**b**), SERBP1 A3240 (**g**), MLLT3 A1068 (**i**) edited by dCasRx-M3. **c**, **h**, **m**, Abundance of ACTB mRNA (**c**), SERBP1 mRNA (**h**), MLLT3 mRNA (**m**) decreased after dCasRx-M3 editing. **d**, mRNA degradation measurement of ACTB in HEK293T cells edited with dCasRx-M3. **e**, mRNA degradation measurement of ACTB in HEK293T cells edited with dCasRx-A5. **i**, **n**, Normalized abundance of altered m6A at SERBP1 A3240 (**i**), MLLT3 A1068 (**n**) edited by dCasRx-A5. **j**, **o**, Abundance of SERBP1 mRNA (**j**), MLLT3 mRNA (**o**) increased after dCasRx-A5 editing. Data is represented as mean ± SEM. (ANOVA, *: P value < 0.05; **: P value < 0.01; ***: P value < 0.001; ****: P value < 0.0001. n = 3).

**Figure S4.**
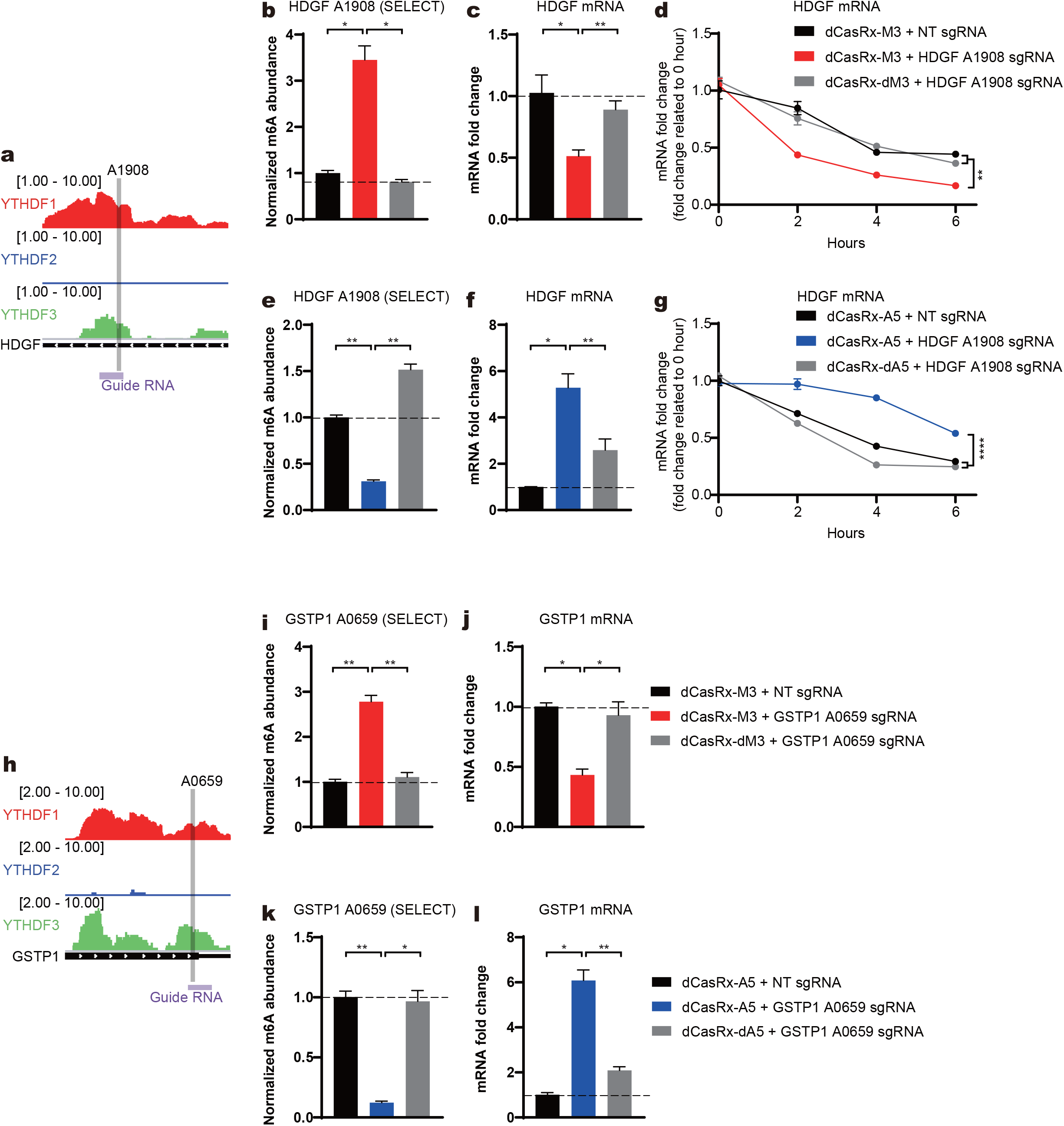
m6A sites binding both DF1 and DF3 control the degradation of endogenous transcripts in HEK293T cell. **a**, **h**, Schematic diagrams of distribution of DF paralogs, DF1 (red), DF2 (blue), and DF3 (green), in endogenous HDGF mRNA (**a**), GSTP1 mRNA (**h**). Grey bars represent dCasRx-M3 or dCasRx-A5 targeted sites. Purple bars represent sgRNA binding location. **b**, **i**, Normalized abundance of altered m6A at HDGF A1908 (**b**), GSTP1 A0859 (**i**) edited by dCasRx-M3. **c**, **j**, Abundance of HDGF mRNA (**c**), GSTP1 mRNA (**j**) decreased after dCasRx-M3 editing. **d**, mRNA degradation measurement of HDGF in HEK293T cells edited with dCasRx-M3. **g**, mRNA degradation measurement of HDGF in HEK293T cells edited with dCasRx-A5. **e**, **k**, Normalized abundance of altered m6A at HDGF A1908 (**e**), GSTP1 A0859 (**k**) edited by dCasRx-A5. **f**, **l**, Abundance of HDGF mRNA (**f**), GSTP1 mRNA (**l**) increased after dCasRx-A5 editing. Data is represented as mean ± SEM. (ANOVA, *: P value < 0.05; **: P value < 0.01; ****: P value < 0.0001. n = 3).

**Supplementary Table S1.**
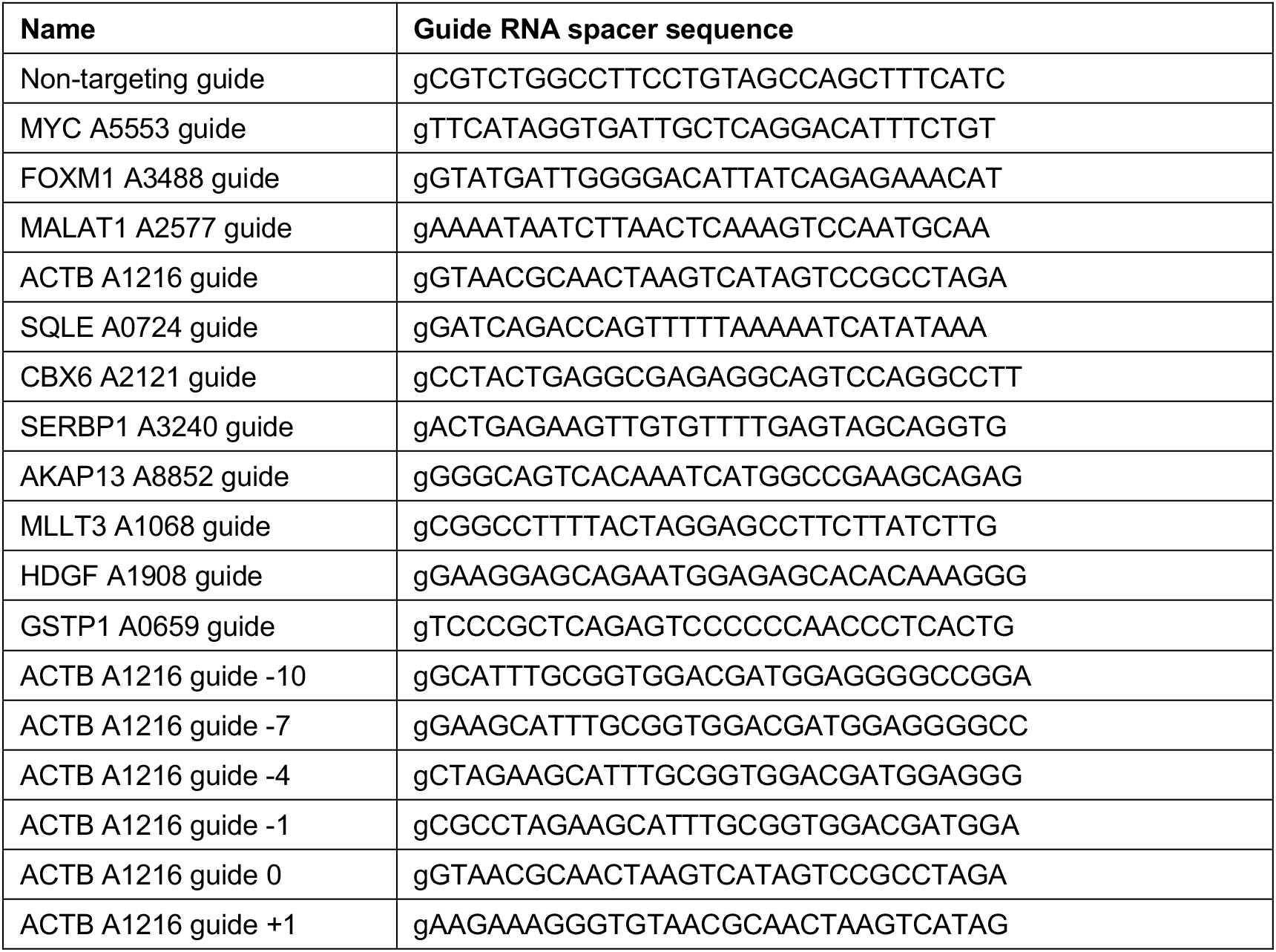
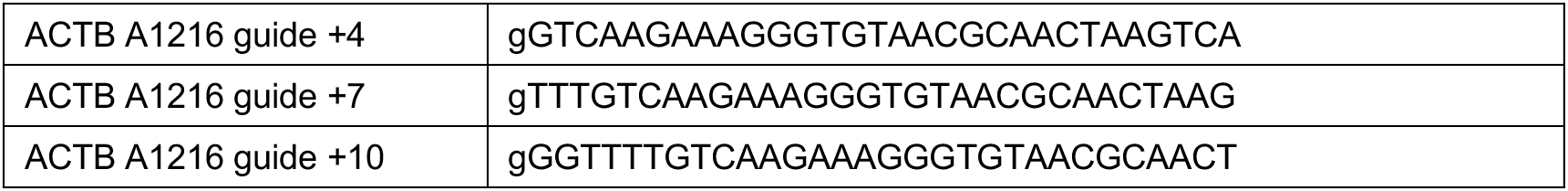
dCasRx sgRNA sequences used in this study.

**Supplementary Table S2.**
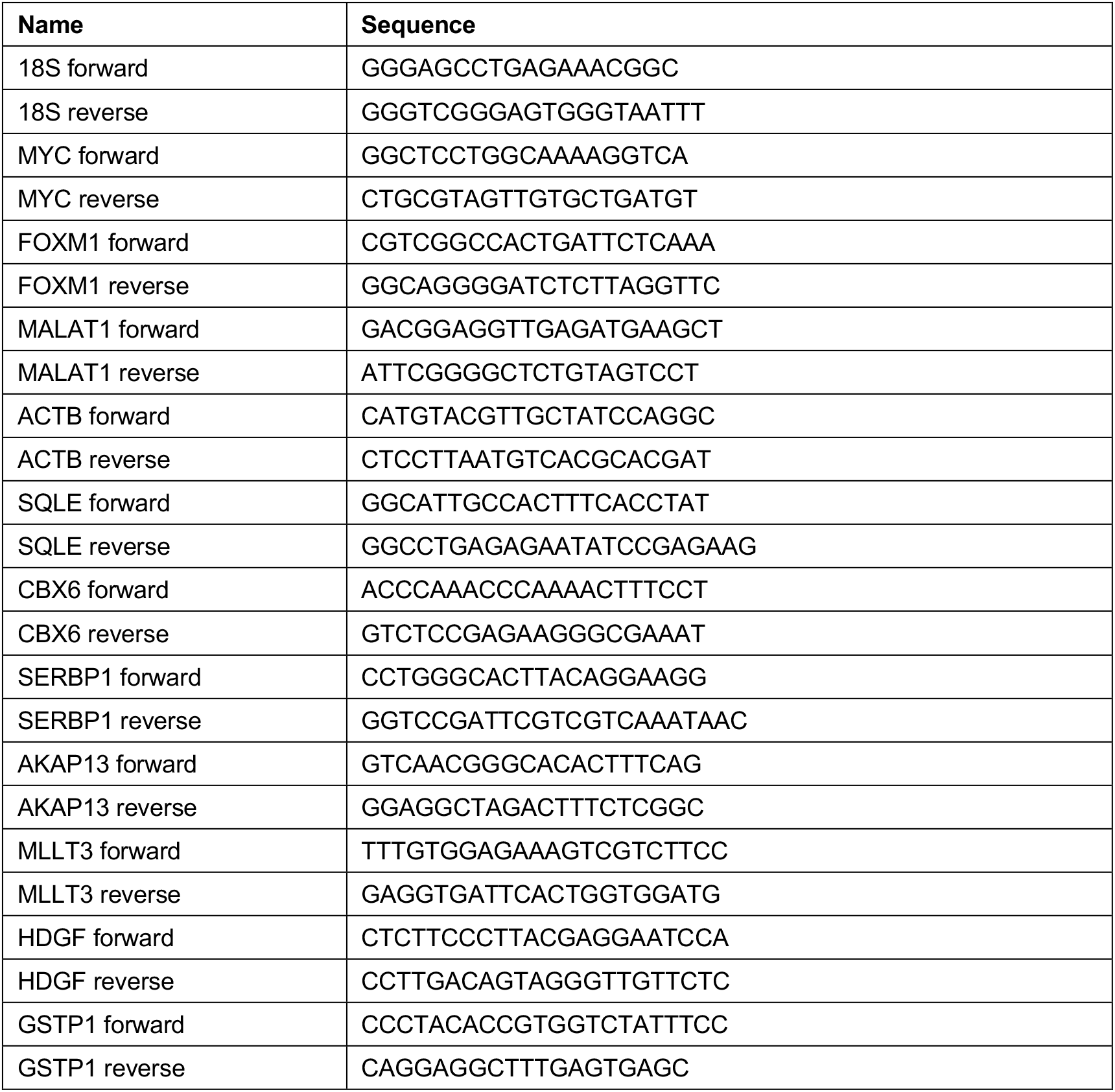
RT-qPCR sequences used in this study.

**Supplementary Table S3.**
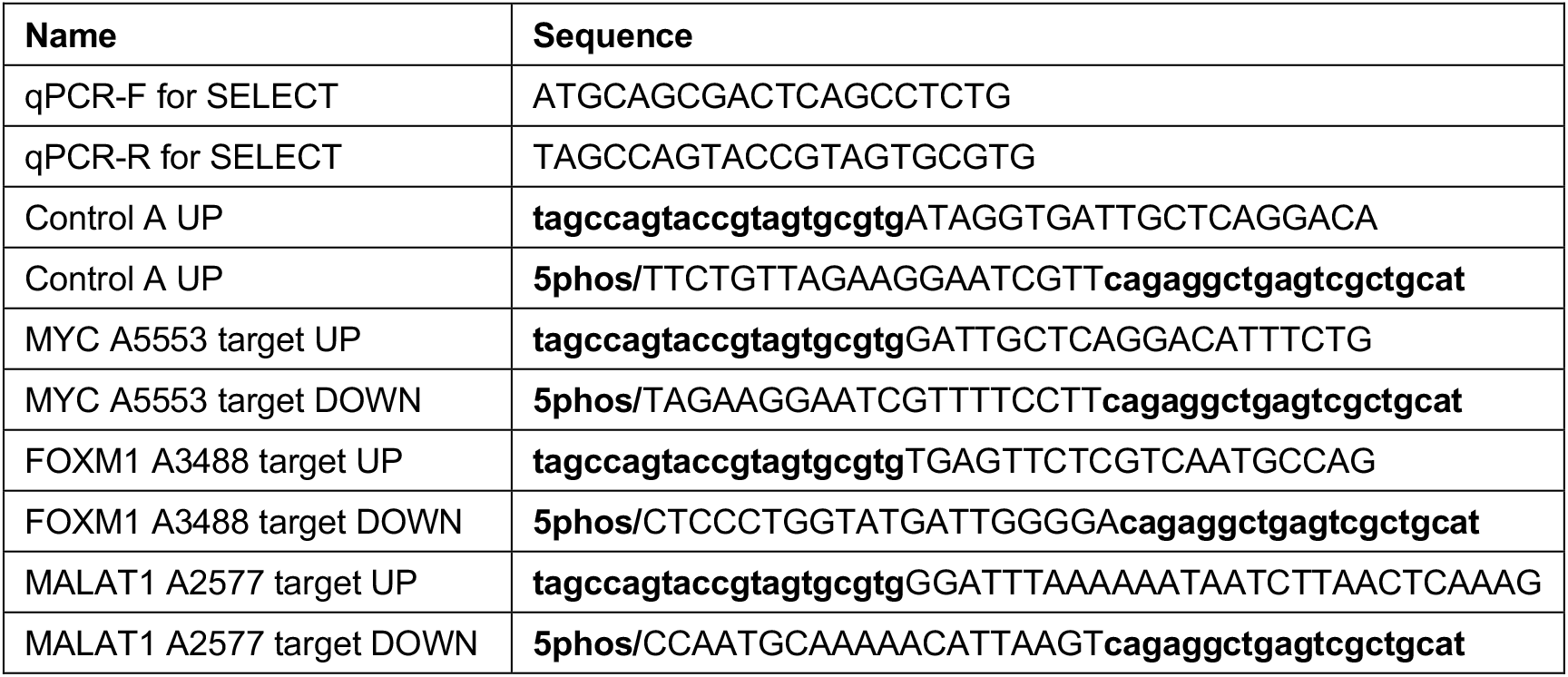
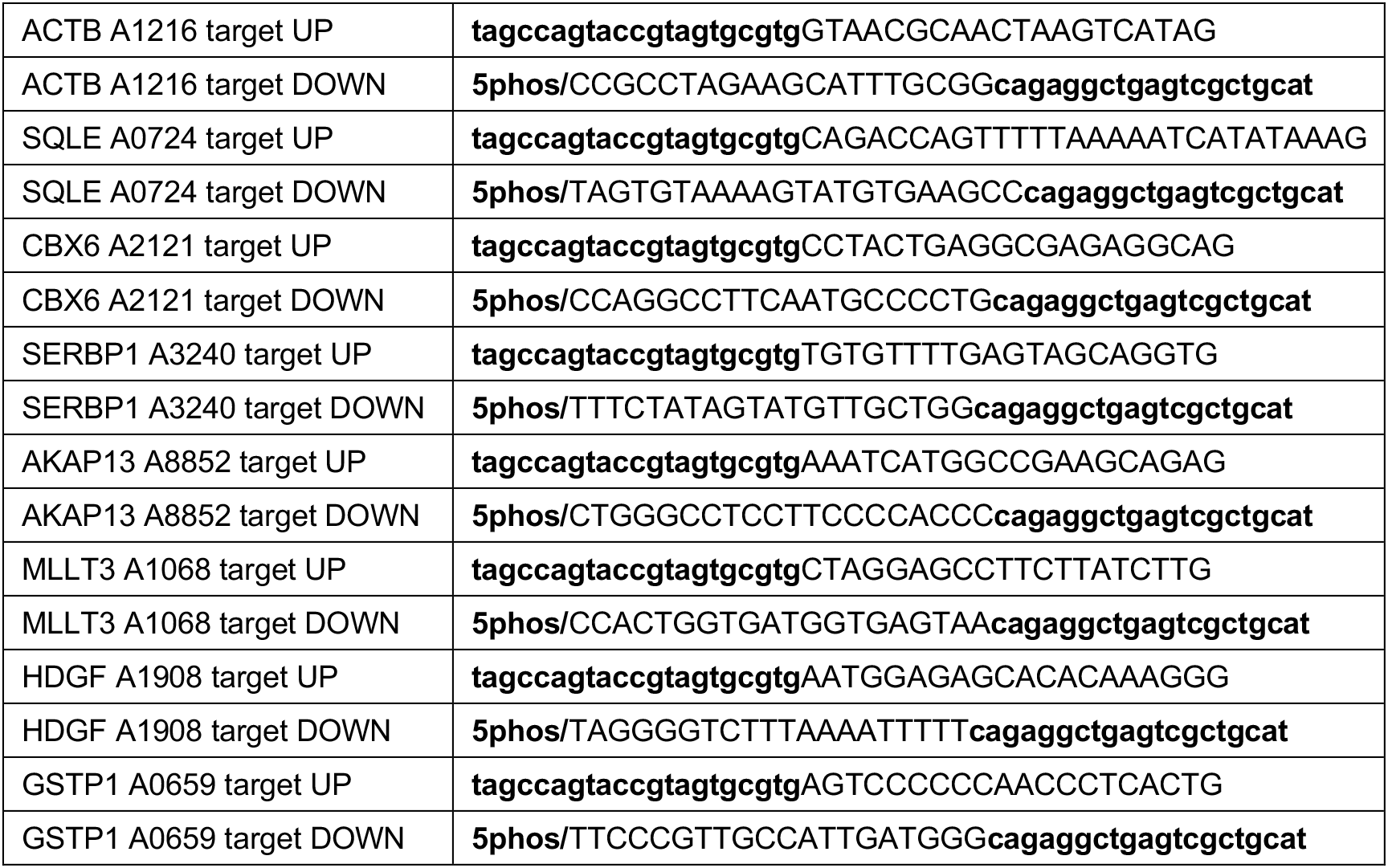
SELECT primer sequences used in this study.

**Supplementary Table S4.**
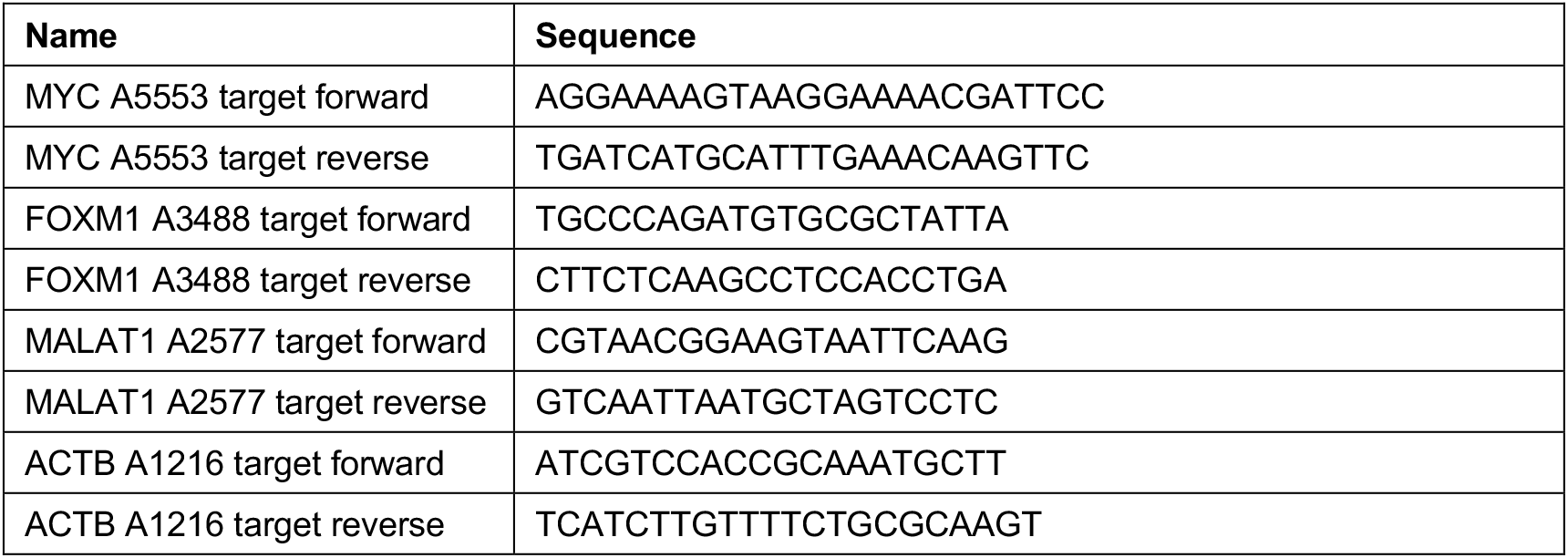
Primers used for m6A-RIP assay in this study.

## Notes

### Competing Interest Statement

The authors have declared no competing interest.

